# ASPEN: A methodology for reconstructing protein evolution with improved accuracy using ensemble models

**DOI:** 10.1101/170787

**Authors:** Roman Sloutsky, Kristen M. Naegle

## Abstract

Evolutionary reconstruction algorithms produce models of the evolutionary history of proteins: the order of duplications and speciations that led to extant homologous proteins observed across species. Although they are regularly used to gain insight into protein function, these models are estimates of an unknowable truth according to the underlying assumptions inherent in each algorithm, its objective function, and the input sequences supplied for reconstruction. In practice, the generated models are highly sensitive to the sequence inputs. In this work, we asked whether we could identify stronger phylogenetic signal by capitalizing on the variance introduced by perturbing the input to evolutionary reconstruction to explore a rich space of possible models that could explain protein evolution. We subsampled from available protein orthologs, “same” proteins across multiple extant species, and produced an ensemble of topologies representing the duplication history which produced related proteins (paralogs) for simulated protein families and in a real protein family – the LacI transcription factor family. We found that two very important phenomena arise from this approach. First, the reproducibility of an all-sequence, single-alignment reconstruction, measured by comparing topologies inferred from 90% subsamples, directly correlates with the accuracy of that single-alignment reconstruction, producing a measurable value for something that has been traditionally unknowable. Second, if we take a large ensemble of trees inferred from 50% subsamples and cast the ensemble into a form that represents the distribution of pairwise leaf distances observed across the ensemble, then trees that capture the most frequently observed relationships are also the most accurate. We propose a new methodology, ASPEN, a meta-algorithm that finds and ranks the trees that are most consistent with observations across the ensemble. Top-ranked ASPEN trees are significantly more accurate than the single-alignment tree produced from all available sequences. Importantly, our findings suggest that the true tree is currently inaccessible for most real protein families. Instead, applications that rely on evolutionary models should integrate across many trees that are equally likely to represent the true evolutionary history of a protein family.

## Introduction

Orthology and paralogy are two forms of evolutionary homology between genetic sequences introduced by Walter Fitch [1, 2] to distinguish between two modes of descent from a common ancestor. Homologs diverged through speciation are called orthologs, while those diverged through the duplication of a genomic region are called paralogs. For coding sequences, paralogs tend to perform related, but distinct functions [3–5]. As a result, families of paralogous proteins provide an excellent opportunity for biochemists and molecular biologists to dissect how functionality is encoded in sequence and structure. Reconstructing histories of paralog divergence can aid tremendously in this endeavor [6–11]. However, such reconstructions can be difficult due to a variety of factors, including, among others: the complexity of the problem, which scales combinatorially with the number of nodes to be reconstructed; the failure of likelihood-based approaches to adequately discriminate between topology models, especially when the amount of phylogenetic data (number of alignment columns) is small (<1000 columns) [12]; and the sensitivity of phylogenetic reconstruction to the input alignment [13–19]. In this work we propose new frameworks for tackling the challenges in reconstruction of protein family divergence, including utilizing model sensitivity to inputs, which is typically thought to be problematic, to improve the accuracy of reconstruction.

To begin, we propose to reduce the complexity of reconstruction, based on the idea that not all ancestral nodes in protein evolution are of equal interest. A typical collection of input sequences comes from multiple species, where the divergence topology for these sequences contains both duplication and speciation ancestral nodes. Depending on the protein family and the number of species represented in the collection, the fraction of speciation nodes may be quite high. Reconstructing speciation history from a single protein family is of little interest aside from identifying extremely rare evolutionary events, such as horizontal gene transfer and reciprocal paralog loss, through reconciliation of the protein phylogeny with the accepted species phylogeny [20, 21]. Unfortunately, the speciation nodes still must be inferred in order to reconstruct any duplication nodes that predate them.

If a high confidence ancestral node can be identified such that no node descended from it is of individual interest – e.g. the ancestor of a set of orthologs diverged exclusively through speciation – then we can imagine “factoring” the topology space search into two components: (a) topologies below the node in question, (b) everything else. This is not particularly helpful for phylogenetic reconstruction in general, since (a) and (b) still must be optimized jointly in order to determine the overall optimal phylogeny. However, if we were not concerned with inferring (a), we would be satisfied to integrate over the uncertainty of (a) to produce, in a sense, a marginal reconstruction of (b). Moreover, treating the root of (a) – the high-confidence ancestral node – as a leaf in our reconstruction of (b), would allow us to consider simultaneously the uncertainty arising from the selection of specific sequences descended from that ancestor to be included in the reconstruction and from their alignment, in addition to the uncertainty in the inference of (a) itself. Here, we propose such an approach, where we attempt to reconstruct the history of protein duplication events, using ortholog sets as defined by known speciation events that gave rise to these orthologs.

Next, we hypothesize that integrating over all sources of uncertainty in the dispensable parts of a protein phylogeny – e.g. ortholog divergence through speciation – would result in better characterization of the phylogenetic signal they contain and improve the accuracy in reconstructing the rest of the phylogeny – duplications which gave rise to paralogs. Testing this hypothesis requires a mechanism for capturing the uncertainty and incorporating it into phylogenetic reconstruction. Inspired by the findings of Salichos and Rokas, who showed that topologies of yeast divergence reconstructed from single genes, while rarely identical, shared much greater similarity than randomly generated topologies [22], we propose a new approach based on the possibility that frequently recurring topological features are more likely to represent phylogenetic signal.

In this work, we present a framework for: 1) assessing the amount of uncertainty arising from selection and alignment of input sequences, 2) identifying topological features and determining the frequency with which they occur across reconstructions, and 3) quantifying the consistency of individual topologies with the identified features. Based on our findings, we propose an observable metric of reconstruction uncertainty, precision, which directly correlates with reconstruction accuracy. We then describe a meta-algorithm that identifies the topologies which are most consistent with topological features extracted from an ensemble of individual reconstructions. Finally, we demonstrate that for some reconstruction tasks, multiple models should be considered as equally likely, given the evidence. Our methodology, ASPEN (Accuracy through Subsampling of Protein EvolutioN), produces significantly more accurate topology models than those produced by a single-alignment reconstruction from all available sequences.

## Results

### Quantifying reconstruction uncertainty with sequence subsampling

#### Defining protein families for testing

We tested the effectiveness of our analysis and inference framework on simulated protein families and the LacI family of bacterial repressor proteins. We turned to simulated sequence data, in addition to a real protein family, for two previously noted reasons [17]. First, simulating evolution over known phylogenies allowed us make a quantitative assessment of reconstruction accuracy compared to the true divergence topology. Second, it allowed us to explore a range of divergence conditions by systematically varying branch lengths of input phylogenies, while controlling for other factors such as overall sequence length and the distribution of secondary structure elements and disordered loops. We simulated 600 protein families, each containing 15 paralogs represented by 66 orthologs – a total of 990 sequences. The scale of this simulation experiment allows for a broad exploration of the effects of topology and divergence time on the accuracy of phylogenetic reconstruction. For simplicity, we used the same tree with 66 species to represent each paralog, resulting in topologies reflecting a series of duplications, producing 15 paralogs in an ancestral genome, followed by speciations leading to 66 divergent genomes, each containing the same 15 paralogs. Evolution of real protein families is rarely that neat. Speciations predating some duplications and loss of some paralogs in some lineages often results in different collections of paralogs appearing in different genomes, dramatically complicating orthology assignments. However, the methodology we describe does not rely on or benefit from this simplification in any way. It only requires the presence of high-confidence ancestral nodes, the topologies below which are of little interest. Since the provenance of collapsed clades does not inform the methodology, the analysis of that methodology based on this simulation experiment is applicable without loss of generality.

**Fig 1.**
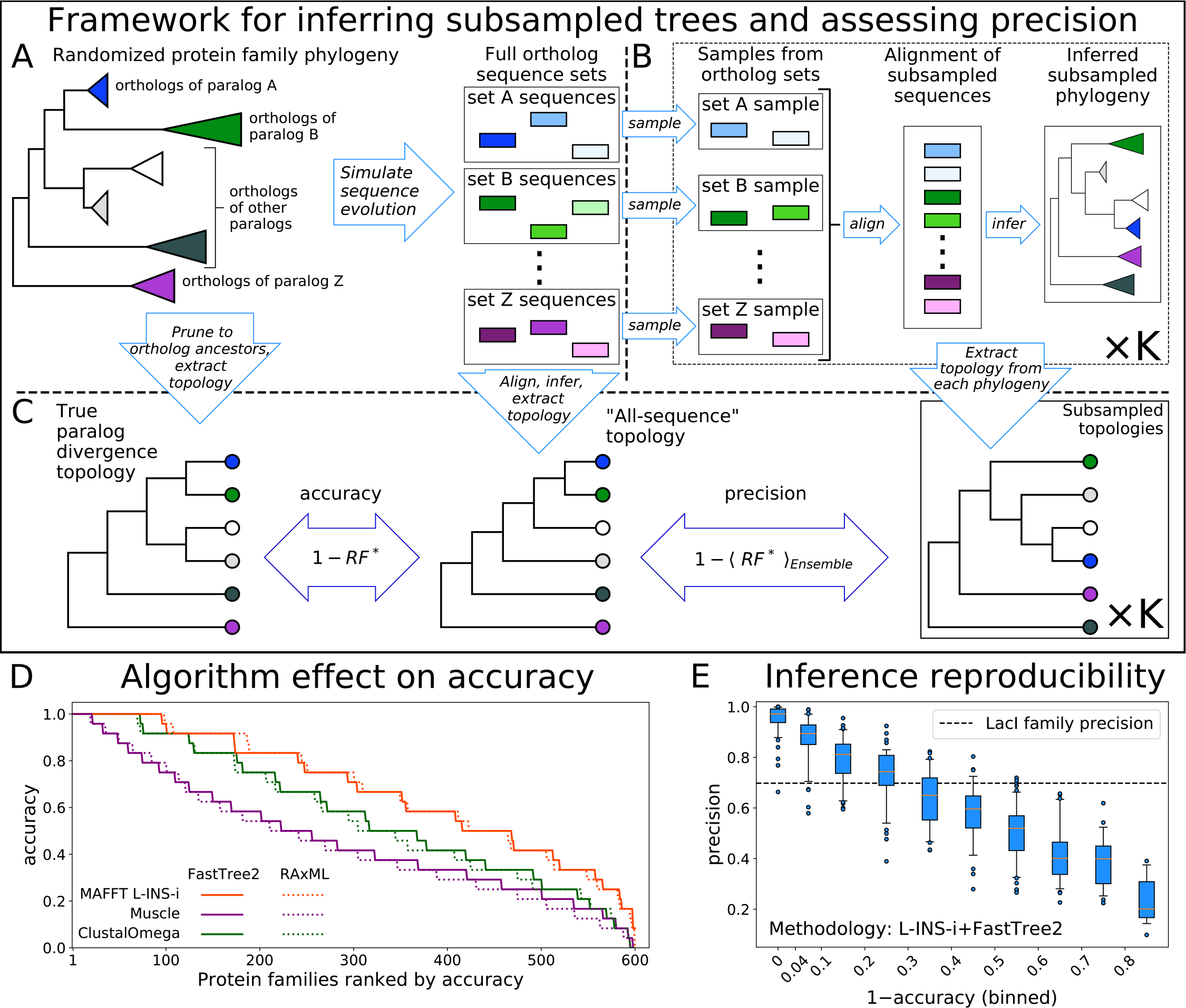
**Top:** Framework for assessing accuracy and precision of reconstruction for simulated protein families using an ensemble of subsampled topologies. (**A**) Sequence evolution was simulated over synthetic phylogenies. (**B**) Sequences were sampled from orthosets without replacement, aligned, and used to infer “subsampled” phylogenies. (**C**) An all-sequence phylogeny was inferred from an alignment of all sequences in a simulated family. The true, all-sequence, and subsampled phylogenies were pruned to orthoset common ancestors. Their branch lengths were discarded to obtain paralog divergence topologies. Modified Robinson-Foulds (*RF*\*) symmetric distance metric was calculated between the true and all-sequence topologies and between the all-sequence and each subsampled topology. Accuracy and precision of reconstruction for a family are defined in terms of these *RF*\* distances. **Bottom:** (**D**) The 600 simulated families are ranked by their all-sequence topology accuracy and plotted according to the alignment and phylogeny inference algorithms used to infer the all-sequence phylogeny. (**E**) Reconstruction precision vs 1 accuracy of the all sequence topology for simulated families. Families are binned by accuracy. Tick marks on x-axis indicate bin boundaries. Dashed line indicates precision of LacI family reconstruction.

#### Framework for generating an ensemble of topologies and quantifying uncertainty

The framework for generating reduced topologies and quantifying reconstruction uncertainty is outlined in Figure 1. Full topologies are reconstructed from sequence data by a combination of alignment and phylogenetic inference methods. We then extract ancestor divergence topologies, with the proposal that branch lengths can be optimized later for any combination of selected topology or collection of topologies and alignment of input sequences. We focus on the divergence of common ancestors of orthologs, dispensing with the divergence of the individual orthologs through speciation. In principle, any ancestor of a subset of input sequences can be used, so long as that ancestor is robustly inferred under reconstruction uncertainty, including uncertainty due to input sequence selection and alignment. In our subsampling framework we define robustness as consistency of ancestor reconstruction across subsamples. In other words, most samples from a set of descendants must be monophyletic in their respective reconstructions in order to consider their common ancestor robustly inferred. We found that subsamples from all of our simulated orthosets were overwhelmingly monophyletic, so our selection of ortholog common ancestors as leaves for our reconstruction was justified.

We quantified topology differences using the Robinson-Foulds symmetric distance metric [23], modified to handle the occasional non-monophyletic reconstruction of an orthoset ancestor (*RF*\*, *Materials and Methods*). When an inferred topology for a synthetic family is compared to the topology over which the family was simulated, that comparison is a measure of the distance between reconstruction and truth. Therefore, we define the accuracy of a reconstruction as:

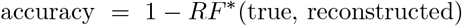

We refer to a topology inferred from an alignment of all available sequences as the “all-sequence” topology. This is the topology most often used when considering a single reconstruction. Using multiple representative sequence sets obtained through subsampling allows us to characterize the uncertainty of ancestor divergence reconstruction arising from selection and, critically, alignment of descendant sequences. In this work we used two different subsampling sizes to characterize uncertainty. First, we subsampled most, but not all, of the available sequences. The comparisons of these topologies to the all-sequence topology is effectively a measure of the reproducibility of the all-sequence reconstruction given highly similar representations of descendant divergence, but under uncertainty due to input sequence alignment. We refer to this quantity as precision, defined as:

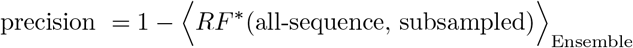

where angle brackets denote average over the ensemble of subsampled topologies. Second, we subsampled roughly half of the sequences from each orthoset to generate larger ensembles, which we used to identify topological features and quantify their consistency with the phylogenetic signal present in the sequence data. Smaller subsamples allow for faster reconstruction of each topology, in turn allowing us to generate larger ensembles, which provide greater resolution of consistency with phylogenetic signal. The distributions of pairwise *RF*\* distances between topologies inferred with 90% and 50% subsampling tend to be centered in the same intermediate range as that observed by Salichos and Rokas for species divergence reconstructions from single yeast genes [22], with the 50% ensemble allowing for a moderately broader exploration of the topology space (S3 Fig). We settled on this sample size as a compromise between strength of phylogenetic signal, breadth of topology space exploration, and inference speed.

#### Selection of alignment and phylogeny inference algorithms

Since the specific alignment and phylogenetic inference algorithms used affect reconstruction [15–18, 24–26], we tested all combinations of three alignment algorithms and two phylogenetic inference algorithms on the 600 simulated families, each containing 990 sequences. We used MAFFT’s L-INS-i protocol [27], ClustalOmega [28], and Muscle [29] for sequence alignment, and FastTree2 [30] and RAxML [31] to infer phylogenies. We compared each all-sequence topology to the true topology and found a wide diversity of accuracies across the 600 synthetic families: accuracy varied between 1, when the all-sequence topology is identical to the true topology, and 0, when *RF*\* between the two topologies is maximal. Therefore, our test set of simulated families spans a range of complexity, which allows us to observe the performance of phylogenetic reconstruction as a function of that complexity. The largest effect on accuracy came from the choice of alignment algorithm, consistent with previous studies [15–18, 24–26]. L-INS-i alignments produced the most accurate reconstructions (Figure 1D). FastTree2 and RAxML performed very similarly across all alignment algorithms, also consistent with earlier observations [26]. Because FastTree2 is significantly faster than RAxML, we selected the fast and accurate combination of L-INS-i and FastTree2 for all subsequent phylogeny inferences.

#### Subsampling produces an observable measure of accuracy

We made a striking observation when we compared the reproducibility of an all-sequence tree with its accuracy. Reproducibility, expressed as precision, was measured by comparing 50 trees created from ≈90% subsamples of available sequences (Figure 1B,C). As shown in Figure 1E, precision directly correlates with accuracy – the most accurate all-sequence trees are also the most reproducible. Precision of reconstruction for the LacI family is 0.698, indicated by a dashed line in Figure 1E, suggesting that the reconstruction is of moderate difficulty and that the all-sequence LacI topology is not entirely accurate, in accordance with its rather low reproducibility. Due to their strong correlation, it is possible that precision – an observable quantity regardless of whether the true topology is known – can be used as a measure of the complexity of reconstruction for a protein family and as a proxy for the accuracy of the all-sequence tree.

We wished to understand why subsampling produces a metric that is correlated with accuracy, a hidden value, and to understand the potential generalizability of this observation. To explore this further, we compared the values of the topology search objective function (log-likelihoods) of topologies, given an alignment:

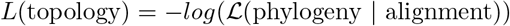

by calculating the fractional log-likelihood difference between a reference topology and an alternate topology according to:

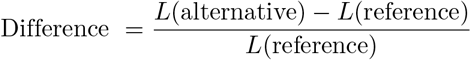

where the reference is the topology selected by FastTree2 for the alignment in question. We used RAxML to evaluate exact likelihood, rather than the approximate likelihood used by FastTree2, because FastTree2 does not provide the functionality to evaluate its objective function without performing a topology search. We systematically compared topologies on alignments using this fractional difference in log-likelihood to effectively measure the distance between models along the likelihood landscape defined by a specific alignment.

**Fig 2.**
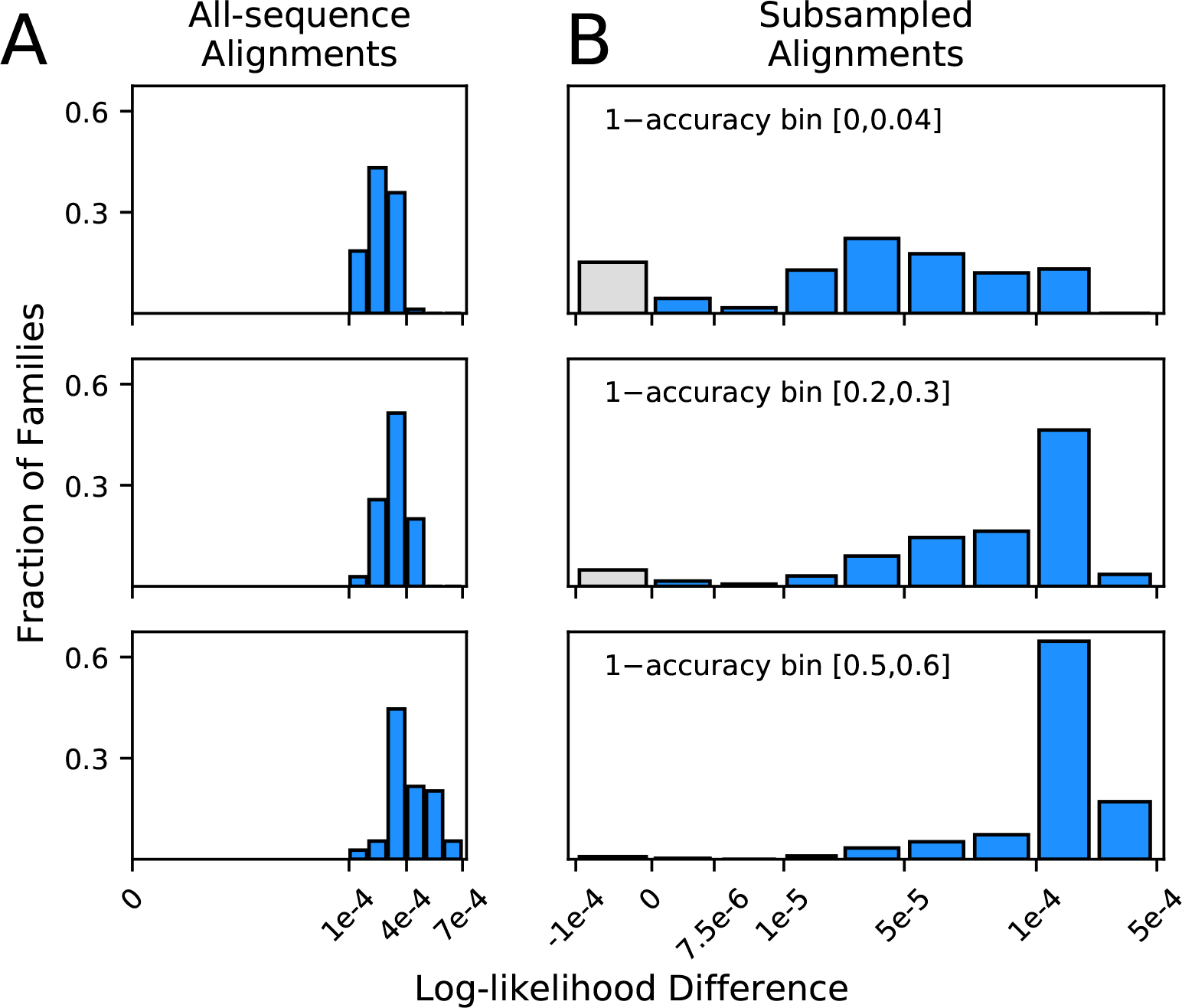
Histograms of log-likelihood differences (calculated using RAxML) between true and all-sequence (reference) topologies, calculated over all-sequence alignments (**A**), and between all-sequence and subsampled (reference) topologies, calculated over subsampled alignments (**B**). Rows present subsampled and all-sequence results for different 1–accuracy bins. Gray bars represent negative difference values in cases where the alternative topology has a higher RAxML likelihood than the topology inferred with FastTree2. These small differences are more likely due to differences between the RAxML and FastTree2 likelihood functions, rather than to topology search failures by FastTree2.

The most immediate question we considered was whether the topology search algorithm simply failed to identify optimal topology models under its own objective function. In particular, we wondered whether this is the reason why the all-sequence alignment failed to recover the true topology for the vast majority of our synthetic families (84.2%). We compared the topologies selected by FastTree2 for all-sequence alignments (the reference topology in our formulation) to true topologies using the log-likelihood difference metric over the all-sequence alignments. A negative value would indicate that the alternate (true) topology obtained a better log-likelihood score than the reference, but the search algorithm failed to identify it. We found no such cases among our synthetic families (Figure 2A). While it is reassuring that topology search reliably identified the better scoring topologies, this finding demonstrates that imperfect accuracy results from the fact that, under the substitution model, the observed differences between input sequences are more likely to have arisen by a sequence of substitutions different from the one that actually occurred when evolution was simulated. Even more surprising, the all-sequence topology scored better both under the substitution model used in phylogenetic inference *and* under the different model used for simulating sequence evolution (S2 Fig, and see Materials and Methods for choice of substitution models). These findings underscore the difficulty of recovering the true topology by phylogenetic inference from short sequences, even using the substitution model under which those sequences were simulated.

Next, we sought to understand how reconstruction of subsampled topologies is related to the accuracy of all-sequence topologies. For a given alignment, the likelihood landscape is a hypothetical multidimensional surface produced by evaluating the likelihood function on topologies, over which the search algorithm seeks the optimal topology for that alignment. We hypothesized that altering the alignment by subsampling sequences and realigning modulates the landscape, relocating the likelihood minima to new topologies, and that the susceptibility of the landscape to such perturbations is related to the accuracy of individual reconstructions. For each subsampled alignment, we compared the subsampled (reference) topology to the all-sequence (alternate) topology using the log-likelihood difference metric. In order to calculate the likelihood of the all-sequence topology given a subsampled alignment, the all-sequence topology was pruned down to the set of leaves contained in the subsampled alignment. Unlike the all-sequence vs. true topology comparisons (Figure 2A), the extent of likelihood differences between subsampled and all-sequence topologies differed considerably with accuracy (Figure 2B). For families with high accuracy, when precision is also high, the difference varied from 0 to 10^−4^, suggesting that the all-sequence topology is within the optimum well. As accuracy and precision decreases, and *RF*\* distances between subsampled and all-sequence topologies increases, the likelihood differences also increases. The subsampling perturbation produces a greater displacement of the optimum, pushing the all-sequence topology out of the optimum well and onto a plateau, where log-likelihood scores are 10^−4^ to 10^−3^, worse than the optimum. Greater displacement of the optimum pushes the all-sequence topology out of the optimum well for more synthetic families, but does not increase their log-likelihood differences from the reference beyond the 10^−4^ – 10^−3^ range. We were surprised to discover that true topologies were always located on this plateau, outside of the optimum well on the all-sequence alignment likelihood surface (Figure 2A). We suggest that the degree of optimum displacement due to resampling and realignment, captured by our precision metric, reflects the overall dispersion of plausible topology models around the true topology. This dispersion of plausible models is the result of each family’s unique divergence history, and both accuracy and precision depend on its extent, explaining why the two are strongly correlated. In other words, less accurate all-sequence topologies are inferred over surfaces more susceptible to the perturbation of input sequence selection and alignment.

### Exploiting reconstruction variance to identify best trees

Next, we sought to apply our sequence subsampling framework to characterize the region of topology space throughout which subsampled topologies are dispersed, and then to use this information to reconstruct topologies that are most consistent with observations across the subsampled ensemble. We arrived at a reconstruction approach that has three parts: 1) a framework to characterize features observed across an ensemble and quantify their occurrence, 2) a scoring function that reflects the consistency of a given topology with observations across the ensemble, and 3) an algorithm that exhaustively enumerates the highest-scoring topologies according to this metric. We call this approach ASPEN, for Accuracy through Subsampling of Protein EvolutioN.

**Fig 3.**
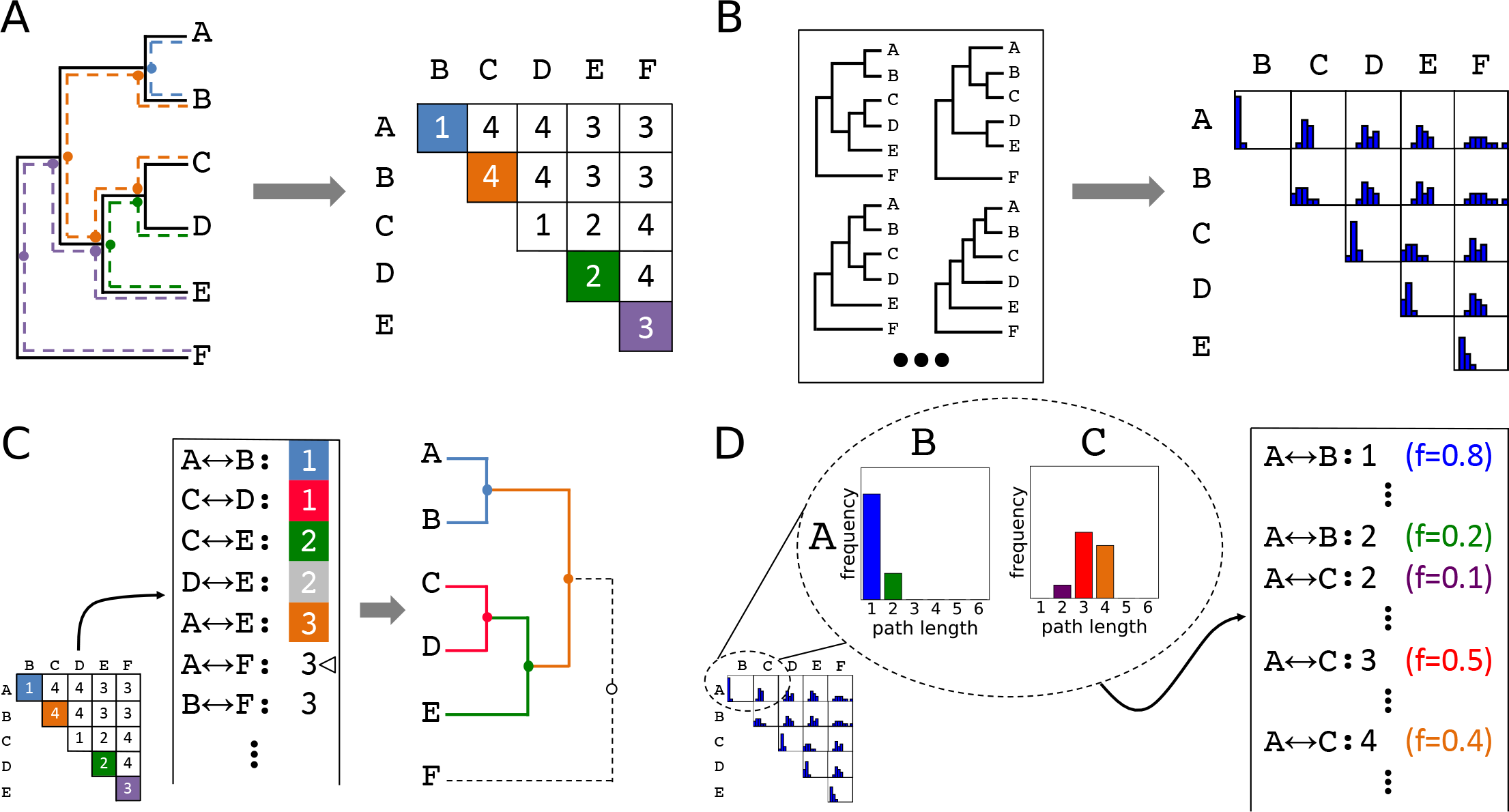
(**A**) Conversion of a topology into a matrix of leaf-to-leaf path lengths. Sample paths (A↔B,1), blue,(D↔E,2), green, (E↔F,3), violet, and (B↔C,4), orange, are highlighted. Dots indicate internal nodes along path. (**B**) Each topology in an ensemble is decomposed into a matrix of leaf-to-leaf path lengths. Observed path lengths for each leaf pair are aggregated into distributions, which are used as weights by ASPEN’s log-frequency scoring function. (**C**) Construction of a topology from its matrix of path lengths representation. First, the matrix is transformed into a sorted list of path lengths. Construction of internal nodes is triggered by path lengths encountered traversing the list. Cursor indicates path (*A↔F*, 3) being recapitulated by construction of node {{{*A, B*}, {{*C, D*}, *E*}}, *F*}. (**D**) The path lengths distribution for each leaf pair produces as many list entries as there were path lengths observed between those leaves. Each path length for each leaf pair has a corresponding observation frequency. These frequencies are used in the scoring function to rank reconstructed topologies.

#### Characterizing topological features across an ensemble

First, we required a way of representing topological features observed across multiple topologies and of quantifying the occurrence of those features. Since there is no obvious way to do this using the standard acyclic graph topology representation, we turned to an alternative representation (Figure 3A) in terms of the number of internal nodes along the paths between every pair of leaf nodes. The two representations are equivalent (Figure 3C), but the pairwise path lengths representation lends itself to aggregating information across topologies in the form of a path length distribution for each leaf pair (Figure 3B), representing empirical probabilities of topological features. Although subsampled reconstructions of simulated orthosets were overwhelmingly monophyletic, occasional reconstructions did contain non-monophyletic ortholog arrangements. This violates an underlying assumption of search space decomposition, as well as the true topology of each simulated family. In such cases, we exclude from the distributions the lengths of any paths from the topology in question that are compromised by passing through spurious nodes resulted from non-monophyletic orthoset reconstruction. Paths from the topology which are not compromised in this way are still included. This highlights a mechanism by which poorly selected ancestors may be identified: when one or more of its putative descendants are consistently placed non-monophyletically with respect to other members of its assigned descendant set, that set’s ancestor cannot be considered high-confidence.

#### Scoring topologies by consistency with identified features

We can use the frequencies with which specific lengths of leaf-to-leaf paths present in a topology occur in an ensemble to reflect the consistency of that topology with observations across the ensemble, and to make comparisons between proposed topologies. ASPEN formalizes this into a scoring function expressed in terms of log-frequencies of leaf-to-leaf path lengths, 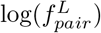, by summing over all pairs of leaves in the topology, according to:

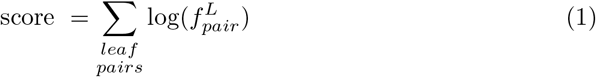

This scoring function rewards incorporation of frequently observed path lengths and penalizes rarely observed ones.

#### Algorithm for constructing *N*-best trees

Given that topologies inferred from all-sequence alignments tend to be inaccurate, increasingly so for more difficult phylogenetic inference problems (Figure 1D and E), ASPEN attempts to identify all likely models of divergence. Specifically, ASPEN’s objective is to identify a set of *N* topologies that are the *N*-most consistent topologies with observations across the ensemble. Given that objective, we created a branch-and-bound procedure to identify the top *N* topologies discussed here. A detailed description of the algorithm is available in Materials and Methods. Briefly, branching occurs when a partially-constructed topology can be extended by multiple internal nodes. An internal node is permitted as an extension only if every pairwise path completed by the proposed node (Figure 3C) appears among the observed path lengths on the list derived from the matrix representation of the subsampled topology ensemble (Figure 3D). Every possible extension is realized in a separate extended topology. Construction of internal nodes is triggered to recapitulate path lengths encountered in traversing the list. Since the number of topologies that might be constructed by this branching can be very large, even given the constraints of ensemble observations, we use bounding to limit construction to the *N* best-scoring topologies. Bounding occurs by checking whether a partially constructed topology might be completed with a better score than the current *N* th-best completed topology. If this is not possible, the partially constructed topology is discarded, bounding all branched construction paths by which it could have been extended. Upon completion of the branch-and-bound procedure, ASPEN will have identified and ranked the *N*-best topologies, according to their consistency with observations from the ensemble of topologies created by subsampling available sequences.

### Evaluation of ASPEN reconstructions

To test the accuracy of ASPEN reconstructions, we used the outlined framework (Figure 1) to generate an ensemble of 1000 subsampled topologies. For this ensemble, we subsampled 30 of 66 orthologs of each paralog in the synthetic families (≈45%) and reconstructed the best 10,000 topologies for 400 families. Since the accuracy of all-sequence topologies varies greatly across synthetic families, and precision is a measure of that accuracy, we binned families by their precision for the purposes of this analysis.

**Fig 4.**
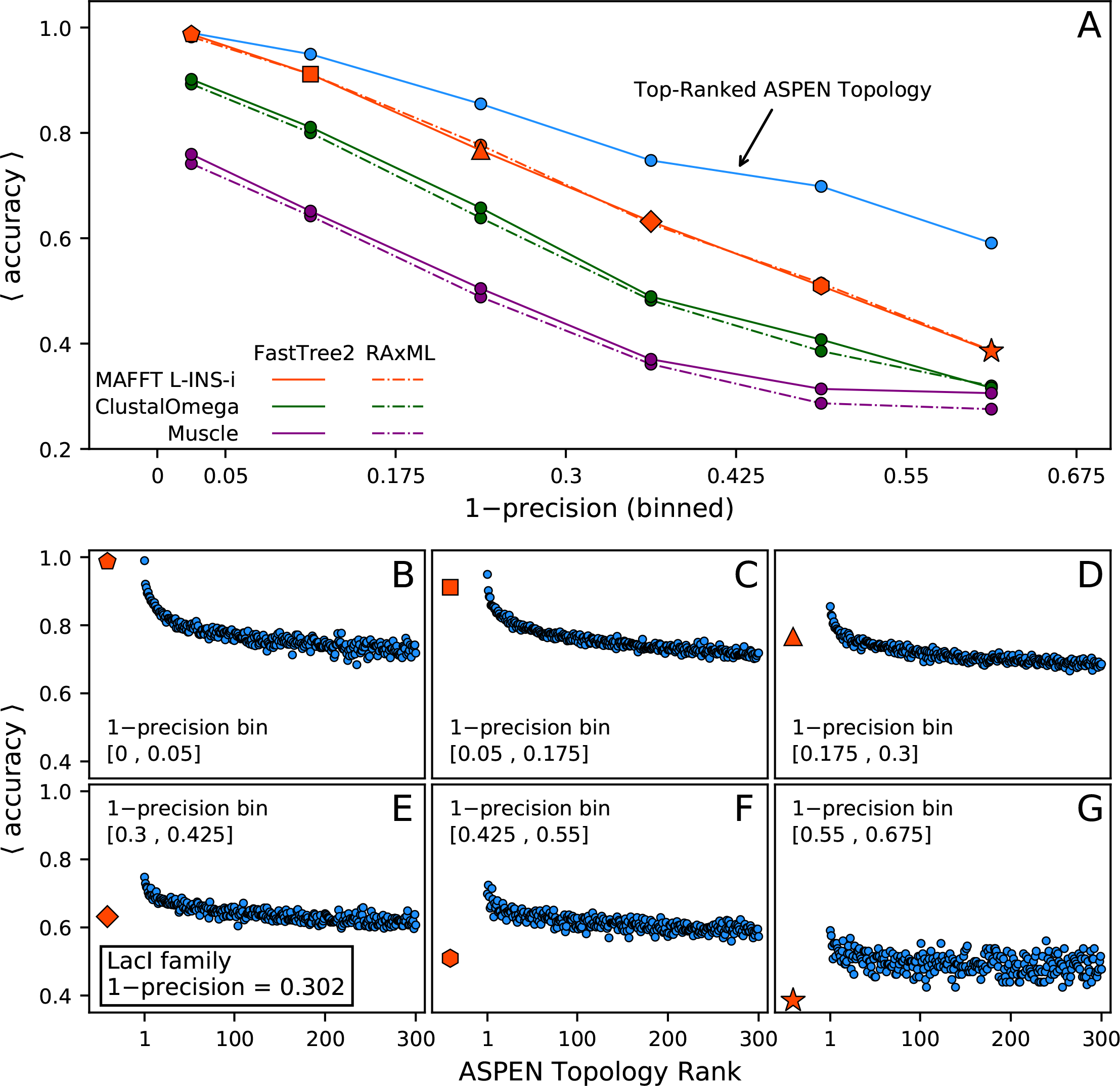
(**A**) Accuracy, as a function of 1–precision, of the top-ranked ASPEN topology and all-sequence reconstructions. Families were binned by precision. Ticks on x-axis correspond to bin edges. Average accuracy across families in bin is plotted for each combination of alignment and phylogeny inference tools. For all-sequence reconstruction with MAFFT L-INS-i and FastTree2 (solid line) a unique marker shape is used in each precision bin. (**B**)-(**G**) For each precision bin in (A), accuracy of ASPEN topologies ranked 1 through 300, averaged within each rank across all families in the bin, is plotted versus rank. Average accuracy of the L-INS-i / FastTree2 all-sequence topologies across the bin is plotted for comparison on the left of each panel.

#### The top-ranked ASPEN topology is the most accurate topology

We compared ASPEN’s top-ranked topology to all-sequence reconstructions using all combination of alignment and phylogeny inference tools (Figure 4A). As with single-alignment approaches, ASPEN’s accuracy correlates with precision, i.e. the complexity of reconstruction for that family. As discussed earlier, MAFFT L-INS-i alignments yielded the most accurate all-sequence reconstructions across all precision bins, while FastTree2 and RAxML performed very similarly on all alignments. Both top-ranked ASPEN topologies and L-INS-i all-sequence reconstructions have nearly perfect accuracy on families in the highest-precision bin. This is not surprising, considering subsampled topology ensembles for ASPEN reconstruction were generated using the combination of L-INS-i and FastTree2. Much more intriguing is the fact that top-ranked ASPEN topologies are consistently more accurate than all-sequence topologies across the remaining precision bins. Moreover, although the accuracy of all reconstructions degrades with complexity of the reconstruction task (lower precision), ASPEN’s accuracy degrades much more slowly. ASPEN’s top topology provides the greatest accuracy improvement over single-topology reconstructions when reconstruction is most complex.

#### Log-frequency score is correlated with accuracy

To understand the relationship between the log-frequency score and the accuracy of reconstructed topologies, we plotted the ASPEN topology rank vs. the bin-average accuracy of topologies (Figure 4B-G). Among higher-precision families (Figure 4B-D), log-frequency scores are strongly correlated with accuracy for topologies ranked in the top ~50. In other words, the score reflecting consistency with ensemble-observed features (Equation 1) is indicative of topology accuracy. The strength of correlation decreases as reconstruction complexity increases (lower precision bins, Figure 4E-G), indicating less discriminatory power with respect to accuracy. Nevertheless, ASPEN’s top-ranked topology is, on average, also its most accurate across all precision bins.

#### ASPEN produces many more accurate topologies

To compare more ASPEN topologies with the most accurate all-sequence topologies, bin-average accuracies of L-INS-i / FastTree2 all-sequence topologies are plotted alongside bin-average accuracies of top-300 ranked ASPEN topologies (Figure 4B-G). Although the log-frequency score provides less discrimination with respect to accuracy, more ASPEN topologies outperform single-alignment topologies as precision decreases. In the two lowest-precision bins (Figure 4F-G), all top-300 ASPEN topologies are more accurate than the most accurate all-sequence topology.

#### How ASPEN produces more accurate topologies

We wanted to understand how log-frequency scoring facilitates identification of more accurate topologies than phylogenetic reconstructions from single alignments. Using the length of path between two leaves, we explored the connection between path length frequencies observed across an ensemble and differences between true, all-sequence, and ASPEN topologies.

**Fig 5.**
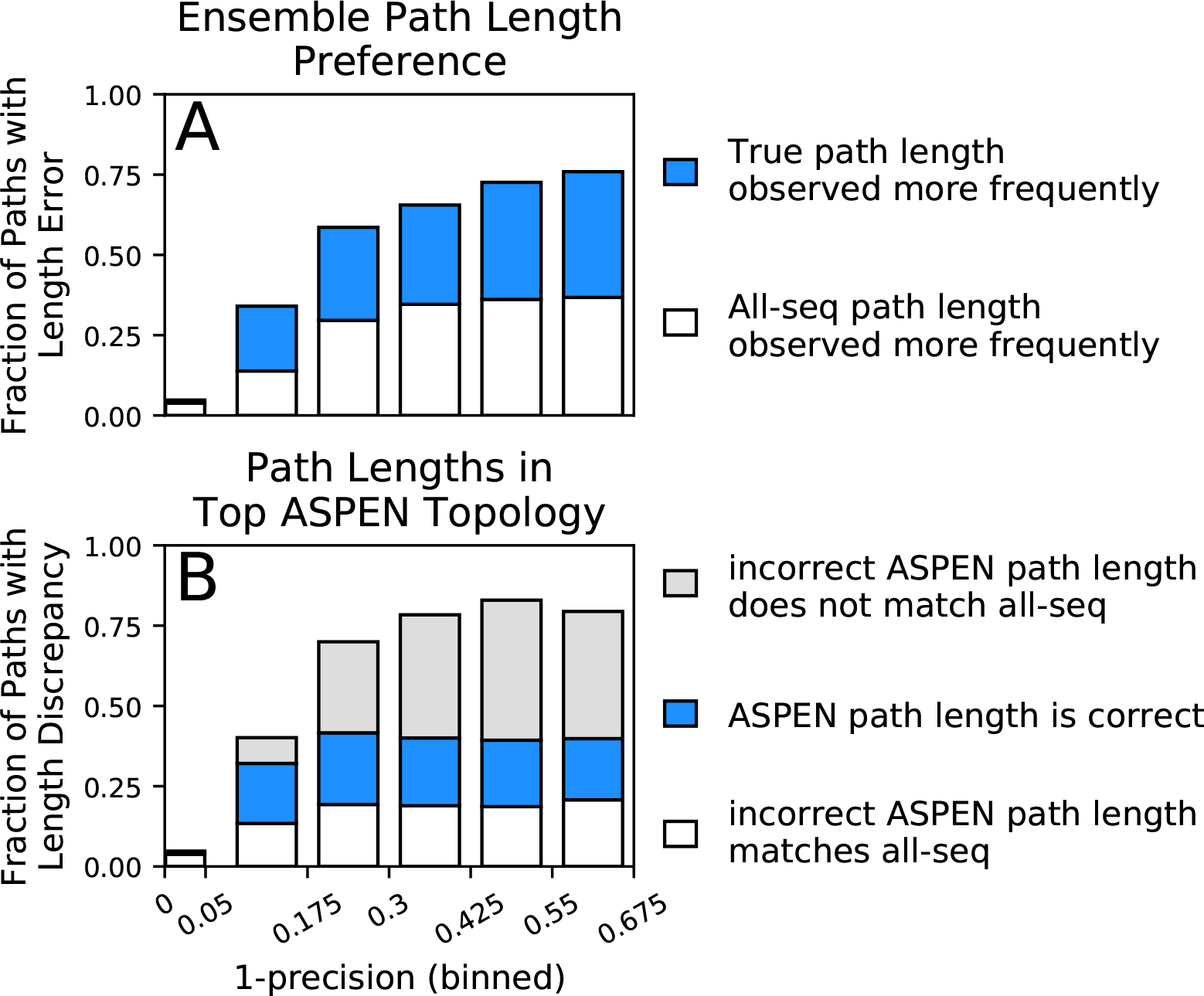
Accuracy of path lengths in the subsampled ensemble and the top ASPEN topology. Synthetic families were binned by 1–precision and path length data was aggregated from all families in a precision bin. (A) Total height of bar represents fraction of all paths across all families in 1–precision bin with a length error (path length is incorrect in all-sequence topology). Hatched fraction of bar represents paths for which the true length was observed more frequently across the ensemble. Empty fraction of bar represents paths for which the incorrect all-sequence length was observed more frequently across the ensemble. (B) Total height of bar represents fraction of all paths in bin with a length discrepancy (on which the true, all-sequence, and top ASPEN topology fail to agree). Hatched fraction of bar represents paths with the correct length in the top ASPEN topology. Empty fraction of bar represents paths on the incorrect length of which the top ASPEN topology agrees with the all-sequence topology. Fraction of bar shaded gray represents paths of unique length in the top ASPEN topologies: incorrect length different from the length in the all-sequence topology.

First, we compared the observation frequencies of path lengths between true and all-sequence topologies. Among paths on which the two topologies disagree, the length consistent with the true topology was observed more frequently than the length consistent with the all-sequence topology in half or more paths across all but the highest-precision bin (Figure 5A). In the highest-precision bin reconstruction is extremely accurate and the true and all-sequence topologies disagree on a very small fraction of path lengths (4.8%). Although the fraction of *all* paths with incorrect length in the all-sequence topology (overall bar height in Figure 5A) increases dramatically as precision falls, the fraction of *disagreeing* paths for which the ensemble correctly identifies the true length (fraction of bar filled with hatched pattern) remains surprisingly constant. The larger fraction of all paths for which the ensemble supports the true path length (height of hatched bar segment) accounts for the much slower drop off in accuracy of all ASPEN topologies at lower precision, compared to the precipitous fall in the accuracy of all-sequence topologies. Unfortunately, the fraction of all paths for which the ensemble supports the incorrect all-sequence path length (height of empty bar segment) also increases at lower precision, which explains why any drop off in ASPEN topology accuracy occurs at all. Nevertheless, the frequency of the most frequent path length and the breadth of the path length distribution provides a measure of confidence in the ensemble’s support of a particular path length. The aggregate of this confidence across all pairwise paths is reflected in the log-frequency score differences between ASPEN topologies.

Next we examined the agreement, in terms of path lengths, between the top ASPEN topology, the true topology, and the all-sequence topology (Figure 5B). As expected, the fraction of all paths with an incorrect length in one or both of the all-sequence and top ASPEN topologies (overall bar height in Figure 5B) is larger at lower precision. Surprisingly, the fractions of all paths with the correct path length in the top ASPEN topology (height of hatched bar segment) and the all-sequence path length in the top ASPEN topology (height of empty bar segment) change little as precision falls, while the fraction of all paths with lengths in the top ASPEN topology matching neither the true length nor the all-sequence length (unique paths, height of gray bar segment) increases. This discrepancy may explain why the accuracy difference between the top ASPEN topology and the other ASPEN topologies decreases at lower precision. Although the fraction of paths with an incorrect length in the all sequence topology, but with the correct length identified through subsampling (height of hatched bar segment in Figure 5A) increases, not all such path lengths are incorporated into the top ASPEN topology – likely due to the constraints imposed by other path lengths on the reconstruction of internal nodes. Instead the correct path lengths are incorporated into other topologies proposed by ASPEN. Accordingly, log-frequency score differences between ASPEN topologies also decrease at lower precision (Figure 6, S4 Fig), reflecting more uniform confidence in any individual topology.

### ASPEN reconstruction of LacI paralog divergence

As mentioned previously, LacI falls into an intermediate range of reconstruction complexity. In this range, the 10 to 30 highest ranked ASPEN topologies are likely to be more accurate than any all-sequence reconstruction, based on our observations from synthetic protein families (Figure 4D,E). Given this, we reconstructed LacI paralog divergence using ASPEN. We derived path length frequencies from an ensemble of 1000 subsampled topologies (Figure 1B), using ~50% of the available ortholog sequences (the same procedure that was used for synthetic families). We then used ASPEN to construct the best 500 topologies (Supplementary Material). Figure 6 plots the drop-off in the log-frequency score of each ASPEN topology, compared to the top-ranked topology, for LacI and for the two 1–precision bins at the boundary of which LacI falls (Figure 4D,E). Log-frequency scores decay faster at higher precision (Figure 6), reflecting a greater difference in confidence for each lower ranked topology, as previously discussed. The all-sequence LacI topology does not appear among the top 500 ASPEN topologies, having scored significantly worse then the ASPEN trees according to the log-frequency scoring function. This indicates that all 500 ASPEN topologies are more consistent with observations across the ensemble of subsampled LacI topologies than the all-sequence reconstruction.

A comparison of all-sequence and top ASPEN topologies (Figure 7) illustrates why the all-sequence topology scores so poorly against the 50% subsampling ensemble. Since the log-frequency scoring function penalizes topologies for incorporating rarely observed leaf-to-leaf path lengths, infrequent incorporation of an ancestral node into ASPEN topologies indicates that most clade arrangements below the node produce unfavorable path lengths. While the top ASPEN topology incorporates the [Mal-B, AscG, GalRS] common ancestor, which appears in and additional 46% of ASPEN topologies, alternative placement of the Mal-B terminal branch in the all-sequence topology produces a different ancestral node, which appears in only 24% of ASPEN topologies. Worse, the all-sequence topology is missing the [CscR, IdnR, GntR] common ancestor, which appears in 76% of ASPEN topologies, incorporating instead the [IdnR, GntR, ExuR, KdgR, FruR, ScrR-BD, TreR] common ancestor, which appears in only 6% of ASPEN topologies. Taken together with our findings for synthetic families, these results suggest that the best ASPEN topology is more accurate than the all-sequence topology, but that none of the reconstructed topologies are likely to match exactly the true divergence of LacI paralogs. In lieu of using a single topology, downstream analyses would do well to reflect this uncertainty by considering multiple likely topologies produced by ASPEN.

**Fig 6.**
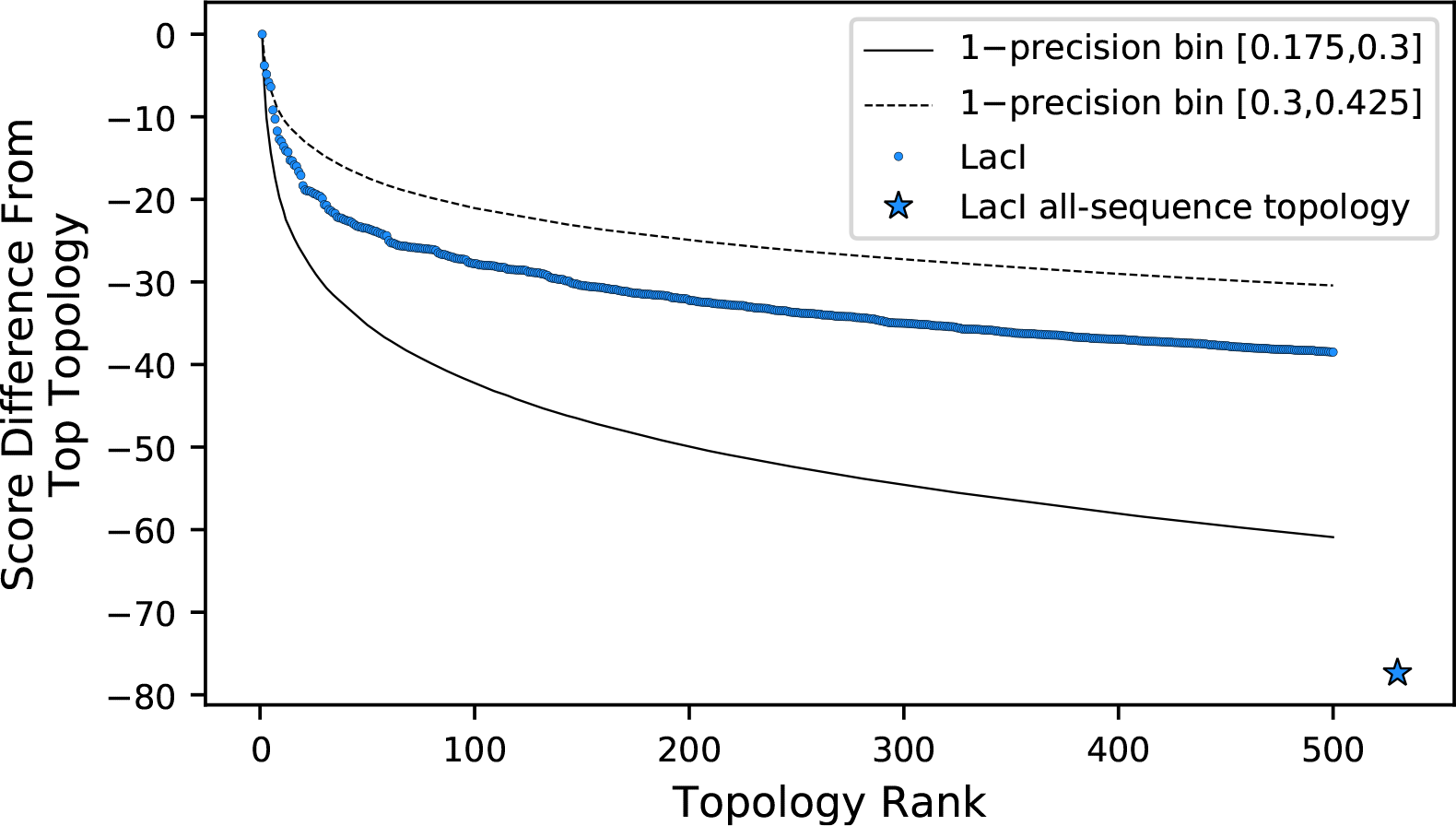
Comparison of log-frequency scores of the top 500 ASPEN topologies for LacI. The difference between a topology’s score and the score of the best ASPEN topology is plotted as a function of topology rank. Bin-average score differences for simulated families from the two precision bins between which LacI falls are plotted for reference. Also plotted is the difference in score between the all-sequence topology of LacI and the top ASPEN topology.

## Discussion

We described a novel approach to analyzing and reconstructing divergence histories of protein families. Our approach is conceptually rooted in the decomposition of the topology search space at high-confidence ancestral nodes, which are extremely likely to exist in the true topology, and takes advantage of the fact that complete divergence histories include “nuisance” segments, which provide little biological insight. Instead of reconstructing such segments, we propose integrating over the uncertainty of their reconstruction to produce more accurate “marginal” reconstructions of the most interesting segments. Critically, our approach considers the uncertainty arising from input sequence selection and alignment, a historically thorny issue in phylogenetic analysis [15]. The traditional method of assessing the reliability of phylogenetic reconstruction, the phylogenetic bootstrap [32], cannot address the reliability of the individual sites (alignment positions) it resamples. On the other hand, the sequence resampling approach we presented occurs farther upstream in the inference process, treating sequence alignment and phylogeny reconstruction as a single inference procedure subject to multiple sources of uncertainty. We can use the resulting ensemble of subsampled topologies 1) to compute an observable metric, precision, which is directly proportional to the accuracy of any individual reconstruction – a hidden quantity for reconstructions from real sequences – and 2) to assemble leaf-to-leaf path length frequency distributions, which we use to define the log-frequency scoring function that is also directly related to reconstruction accuracy. Our topology reconstruction algorithm then uses the scoring function to identify and rank topologies according to their consistency with the phylogenetic signal characterized by these empirical distributions. The highest scoring topologies are more accurate than topologies reconstructed from alignments of all available sequences, confirming that the topological features more frequently represented across subsampled topologies are also more consistent with the true phylogenetic signal. Crucially, ASPEN identifies these topologies in the face of misleading likelihood landscapes resulting from each individual input alignment, on which the true topology is not the maximum likelihood topology, or even located within the optimum well. Finally, we showed that, although ASPEN tree accuracy declines as the reconstruction task gets harder (as evidenced by decreased precision and accuracy of the all-sequence tree), its decline is significantly slower than that of single all-sequence reconstructions. Importantly, ASPEN is able to identify when an all-sequence tree is likely to be inaccurate (via precision) and then construct and rank a set of trees which, while unlikely to be exactly correct, are all likely to be more accurate than any all-sequence tree. We propose that downstream analysis relying on a divergence topology should aim to integrate over this topological uncertainty.

**Fig 7.**
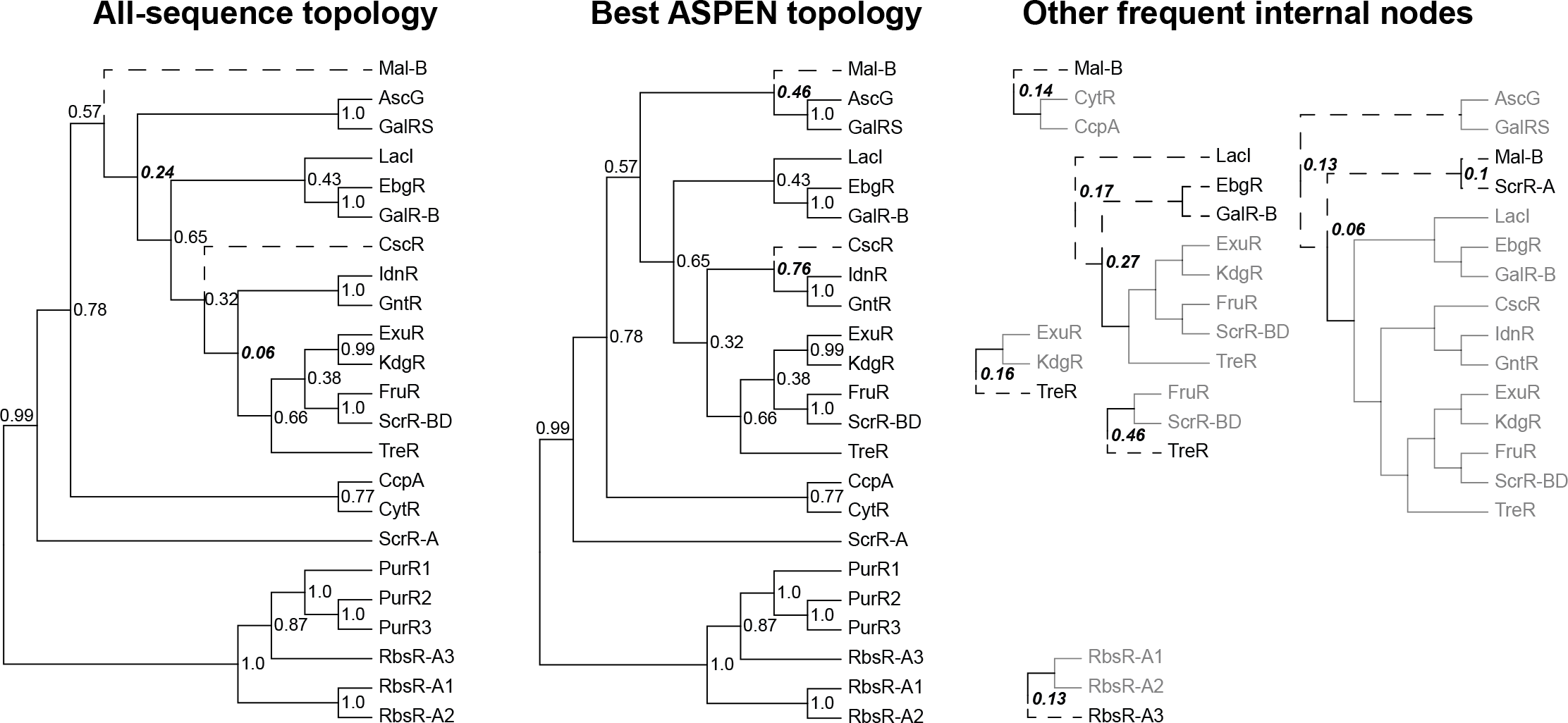
Reconstructed topologies for the LacI family. Reconstructed nodes are annotated with the frequencies at which they were recapitulated among the 500 top-scoring topologies reconstructed by ASPEN as a way of summarizing ancestral nodes observed across the most likely trees. Subtrees on right represent reconstructed nodes observed with frequency ≥0.1 among the 500 ASPEN topologies, but not appearing in either the all-sequence or the best ASPEN topology. Branches placed differently in the all-sequence and best ASPEN topologies are shown as dashed lines, as are branches placed differently from either topology in the subtrees on the right. Observation frequencies for disagreeing splits are bolded and italicized.

Divergence reconstructions for well-studied protein domain families are relied upon extensively by the scientific community. For example, evolutionary trees of catalytic and recognition protein domains involved in signaling, including protein kinases [33] and phosphatases [34], SH2 domains [35], de-ubiquitinating enzymes (deubiquitinases or DUBs) [36], histone deacetylases (HDACs) [37], and Ras GTPases [38] are ubiquitously used. Because such reconstructions are created from single sequence alignments, they ignore the great deal of uncertainty in topology reconstruction under equally valid representations of available sequence data. Furthermore, such reconstructions are often built from limited data. For example the human kinome, which has been cited over 6,000 times to date, was constructed just from human sequences – an example of extreme subsampling, with each ancestor represented by a single sequence. Topology reconstructions from single alignments with sparse subsampling are likely to be even less accurate. We found individual reconstructions are extremely unreliable, even for relatively high-precision families, with very few descendants representing each ancestor. These observations suggest that revisiting these important protein domains using ASPEN’s approach, including quantifying the likely accuracy of published trees and constructing and ranking trees most consistent with the available homolog sequences, is worthwhile. Specifically, we propose that for most protein families, it will be necessary to consider multiple equally likely models of evolutionary divergence.

### Practical considerations in applying ASPEN approach

The methodology we presented can be used to reconstruct the divergence of ancestors in real protein families more accurately than single alignment phylogenetic inference. Selecting sequence sets for subsampling, thereby designating their common ancestors as sites of search space decomposition, is the first step in any analysis. While common ancestors of orthologs are natural candidates when they can be clearly identified, the only requirement is sufficient confidence in the ancestor’s existence. The simulation scheme we used to generate synthetic data used the same species tree for the divergence of each orthoset, producing phylogenies with easily identifiable orthosets, but the ortholog ancestors we selected for the analysis could be identified *de novo* based on their reproducibility across the subsampled ensembles. In our analysis of the LacI family we used this criterion, together with genomic annotations, to select the ancestral nodes (see Materials and Methods). The sequence sets we selected did not come from a uniform collection of species, indicating that post-speciation duplications and gene loss occurred in the evolution of LacI paralogs. Although this step must be carried out individually for any family based on the information researchers wish to obtain from the analysis, we can provide some suggestions for how to apply our methodology to one’s protein family of interest.

We recommend relying on existing genomic and/or functional annotations and average in-group vs. average out-of-group sequence similarity (separability) in designating sequence subsets. When dealing with large families, such as some protein domains, which can number in the dozens or hundreds in vertebrate genomes, common ancestors of multiple paralogs may also be logical choices. The methodology can be used recursively to reconstruct the divergence of these paralogs separately – another advantage of decomposing the search space at high confidence nodes. Similarly, collections containing sequences of uncertain provenance can be analyzed separately and integrated into a larger phylogeny using our approach. Next, we recommend reconstructing an all-sequence topology and a small ensemble of densely subsampled topologies to determine the reconstruction precision and form an expectation of the ultimate accuracy of reconstruction. If selected sequence subsets are not reliably monophyletic across these topologies, the selections need to be revisited until the common ancestor of each subset can be inferred with high confidence. Poorly behaved sequences, those that jump between sequence sets from reconstruction to reconstruction, can either: a) be treated as single representatives of an ancestor and “resampled” (included) in every subsampled topology, or b) withheld from subsequent analysis and grafted later using phylogenetic placement. Once sequence subset assignments are finalized and precision has been assessed, researchers can proceed with the construction of a larger subsampled ensemble, tabulation of empirical path length distributions, and ASPEN topology reconstruction.

ASPEN’s branch-and-bound algorithm provides a powerful guarantee of completeness – that the *N*-best trees were produced – at the end of its run, but execution times and resource requirements can be substantial for large families. However, the vast majority of *N*-best trees are identified very early in the the run, with the remainder of the run spent almost exclusively rejecting worse topologies. Dispensing with the branch-and-bound guarantee, topology assembly can be truncated after a small fraction of the full run time, retaining a nearly-complete collection of *N*-best trees.

Once ASPEN has produced a collection of likely topologies of ancestor divergence, researchers may want to obtain complete phylogenies for their input sequences. In cases where ortholog ancestors were used, we recommend using the species divergence topology for the divergence of orthologs, unless truly compelling evidence to the contrary exists. In our analysis of simulated families, where ortholog evolution was simulated over the same species tree in each case, the correct speciation topology was never recovered for all 15 paralogs, underscoring the futility of reconstructing species divergence from single protein families. Once complete topologies have been assembled from ASPEN topologies and species trees, branch lengths and other parameters can be optimized for any given sequence alignment. Difficult sequences can be attached at this time by phylogenetic placement. Resulting phylogenies can then be used for downstream analyses.

### Extensions to the methodology

We anticipate that, as a meta analysis approach to tree evaluation and reconstruction, ASPEN is likely to continue to boost the accuracy of individual alignment and tree reconstruction approaches, regardless of the specific underlying alignment and reconstruction algorithms. Alternate statistical approaches are increasingly important with the advent of affordable genome sequencing and the resulting explosion in the number of sequenced and annotated species’ genomes [39, 40]. Our entire methodology scales much better with the total number of input sequences than traditional phylogenetic approaches due to the decomposition of the topology search space, although further studies are necessary to explore the effects of tree size on the relationship between precision and accuracy and the signal to noise across an ensemble. The current instantiation of ASPEN as a subsampling and scoring approach is immediately tractable for large protein families. ASPEN’s branch-and-bound reconstruction algorithm is also immediately tractable for reasonably sized families, such as the LacI family. However, the sequence subsampling approach, the path-length frequency distributions it provides, and the log-frequency scoring function are powerful tools in their own right, which scale with the number of *sequence sets* (selected ancestors) much better than the branch-and-bound algorithm. A more efficient search of topology space under this objective function, with a completeness guarantee and/or estimation, is possible. The impact of the subsampling fraction and other aspects of the subsampling methodology on both accuracy and speed in topology scoring and reconstruction also warrants further consideration. It may be possible to structure subsampling so that the resulting frequency distributions are even more consistent with the true topology. Finally, a mechanism for robust inference of branch lengths for ASPEN-constructed topologies that, similar to ASPEN topology reconstruction, integrates over the uncertainty of alignment and reconstruction below selected ancestors, is clearly desirable.

## Materials and methods

### Sequences

#### Simulated paralog families

We simulated 600 families, each containing 15 paralogs, with each paralog represented by 66 orthologs. First, we generated random 15-leaf phylogenies representing paralog divergence. Random phylogenies were generated with average branch lengths of 0.5, 0.6, 0.7, 0.8, 0.9, and 1.0 – 100 phylogenies each. Next, the Ensembl Compara species tree topology [41] containing 66 metazoan species was grafted to each leaf of each random topology to represent ortholog divergence. Finally, each species tree topology was parametrized with branch lengths corresponding to species divergence times obtained from timetree.org [42, 43], randomly rescaled in total height to represent faster or slower evolution of individual paralogs, and then had each individual segment randomly perturbed around its previous length. Sequence evolution was simulated over each resulting phylogeny, seeded with an alignment of human tyrosine kinase domains with median length of 269 a.a. All sequence simulation materials and simulated sequence alignments are available via Figshare (10.6084/m9.figshare.5263885).

#### LacI transcription factor family

We started with a collection of 19 LacI paralogs represented by 28 to 192 orthologs [44]. After initial phylogenetic reconstruction we split paralogs PurR and RbsR-A into three separate paralogs each, according to monophyletic grouping of orthologs. This resulted in new paralogs PurR1 (37 orthologs), PurR2 (61 orthologs), PurR3 (28 orthologs), RbsR-A1 (79 orthologs), RbsR-A2 (22 orthologs), and RbsR-A3 (45 orthologs). The final collection contains 23 LacI paralogs represented by 22 to 192 orthologs, for a total of 1777 sequences (Supplementary Material).

### ASPEN topology reconstruction algorithm

#### Equivalence of topology representations

We demonstrate equivalence of acyclic graph and path length matrix representations of individual topologies by presenting a procedure for interconverting between the two. Transformation of a topology into its path lengths matrix representation is trivially accomplished by counting internal nodes along each path between pairs of leaves (Figure 3A). The reverse transformation can be accomplished using a simple bottom-up construction procedure. Figure 3C provides an illustration by reconstructing the topology from Figure 3A starting with its matrix representation. Because all path lengths are derived from a single topology, they are guaranteed to be consistent, making the construction unambiguous.

#### A branch-and-bound topology construction algorithm

Using the bottom-up procedure for reconstructing a single topology graph as a template, we developed an algorithm that uses a branch-and-bound strategy to construct the requested number of highest-scoring topologies according to the log-frequency scoring function (Eq. 1). By analogy with the single-topology procedure, path lengths, together with their observation frequencies, are sorted into a list (Figure 3D). This list guides topology reconstruction (Figure 3C). However, unlike the single-topology case, list entries are not necessarily consistent with each other. The simplest illustration of this are paths of different lengths between the same two leaves, Figure 3D: e.g. (*A↔B*, 1) observed in the ensemble 80% of the time and (*A↔B*, 2) observed 20% of the time. Such paths are clearly mutually exclusive. The branching component of branch-and-bound accommodates the divergent topologies which recapitulate each path.

##### Branching

As in the single-topology procedure, construction of internal nodes is triggered by path length entries encountered during list traversal, with one key difference. In single topology reconstruction, if a path length could be recapitulated by the introduction of an internal node, that node could be safely constructed because it was guaranteed to satisfy every other list entry. Since that guarantee no longer holds, multiple topologies are constructed simultaneously by allowing the construction sequence to branch (S1 Fig). “Assemblies” are used to track simultaneous reconstruction of multiple topologies. Each assembly holds a copy of the path length frequencies list, a partially constructed topology, and the current topology score according to the scoring function. Reconstruction proceeds in iterations, starting with a single empty assembly on the first iteration. The entire list is traversed and every possible extension is created simultaneously in a copy of the original assembly. In each resulting assembly, all path lengths completed by the new node and all path lengths incompatible with it are marked and not re-examined on subsequent iterations. Remaining path lengths are not completed by the new node, but remain compatible with it. On subsequent iterations the same procedure is repeated for all tracked assemblies.

In principle, branching and iteration alone yield every topology consistent with path lengths observed in the ensemble. In practice, this results in a combinatorial explosion of tracked assemblies, which must be carefully managed to allow construction to proceed to completion.

First, branching to satisfy non-conflicting path lengths can lead to collisions between diverged construction sequences on later iterations (S1 Fig). This occurs because most topologies can be constructed by introducing internal nodes in multiple orders. Each branched construction sequence represents a particular order of internal node introduction. In a practical implementation these collisions must be managed in order to prevent construction of the same topology by multiple construction sequences – an enormous replication of effort.

Second, even if each distinct topology is constructed once, in most cases reconstructing every topology consistent with observations from the ensemble, no matter how infrequent, is neither practical nor useful. Bounding, described in the next section, guarantees reconstruction of only the requested number of top-scoring topologies.

##### Bounding

Completed topologies are ranked according to their log-frequency score, with the ranking updated every time a new topology is finalized. The number of top scoring topologies to reconstruct, *N*, is specified at the beginning of a reconstruction run. Once the initial *N* topologies have been constructed, the *N* th topology score constitutes the bound. Partially constructed topologies (assemblies) are abandoned if no complete topology can be derived from their construction state with a score above the current bound. We determine this by calculating the score for already-incorporated path lengths and projecting the best possible score for a complete topology by assuming the most frequent remaining path length will be incorporated for every unconnected leaf pair:

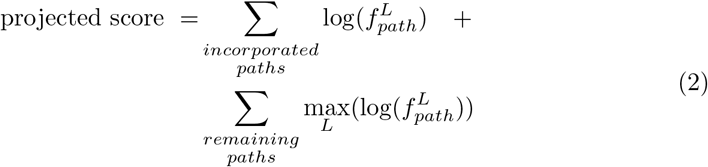

As more high-scoring topologies are constructed, the bounding criterion becomes more strict, allowing both more and earlier abandoning of unproductive construction sequences. The branch-and-bound strategy guarantees that the *N* topologies remaining on the list after all active assemblies have been completed or abandoned are the *N* highest scoring topologies according to the scoring function.

### Simulation of sequence evolution

Random 15-leaf phylogenies were generated at www.trex.uqam.ca [45] using the procedure of Kuhner and Felsenstein [46]. Human tyrosine kinase domains were aligned using MAFFT L-INS-i with default parameters. This alignment was used as the template for sequence simulations as follows. The alignment was divided into 24 segments on the basis of local sequence similarity and analysis of solved tyrosine kinase structures. Each segment was assigned a substitution rate scaling factor and an insertion/deletion model to match degree of conservation and solvent exposure in solved structures. Simulation was carried out over synthetic phylogenies using indel-seq-gen [47–49] under the JTT substitution model.

All sequence simulation materials, including synthetic phylogenies, the template alignment, and indel-seq-gen control files, as well as simulated sequence alignments are available via Figshare (10.6084/m9.figshare.5263885).

### Phylogeny inference

All-sequence phylogenies were inferred using all combinations of MAFFT L-INS-i, ClustalOmega, and Muscle for sequence alignment with FastTree2 and RAxML for phylogeny inference. Subsampled phylogenies for precision calculations were inferred with FastTree2 only, due to run time considerations. Subsampled phylogenies for ensembles used by ASPEN were reconstructed using L-INS-i and FastTree2 only.

Alignment algorithms were used with their default settings. FastTree2 was used with default settings and the WAG substitution model. RAxML was used with default settings and the PROTGAMMAWAGF variant of the WAG substitution model. The WAG substitution model was deliberately used for topology inference, instead of the JTT substitution model used for simulating protein families, in order to emulate the more realistic scenario where models used for reconstruction of phylogenies for natural families do not precisely match the substitution patterns in those families.

Accuracy and precision of reconstruction for a protein family are defined in terms of the L-INS-i / FastTree2 all-sequence and subsampled topologies.

### Modified Robinson-Foulds topology distance metric

The Robinson-Foulds [23] (*RF*) metric is defined in terms of leaf partitions at internal topology nodes for two topologies with identical sets of leaves. For a tree with *N* leaves there are *N* − 3 informative splits. The normalized form of the Robinson-Foulds comparison metric for two topologies, *A* and *B*, is:

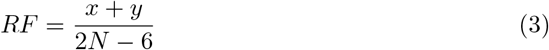

Where *x* is the number of leaf partitions in *A* but not in *B*, *y* is the number of leaf partitions in *B* but not in *A*, *N* is the number of leaves in each topology, and 2*N* − 6 = 2 × (*N* − 3) is the total number of informative splits in the two topologies.

In order to compare reconstructed ancestor divergence topologies we had to modify the *RF* metric to accommodate cases when the ancestor of a descendant sequence set has as descendants one or more other ancestors (non-monophyletic reconstruction). Such topologies are poorly formed because they require inference of additional unobservable events – loss of paralogs in some lineages – in order to be reconciled with a duplication/speciation divergence history. Because the offending subsample cannot be pruned to a common ancestor leaf, the resulting topology cannot be compared to properly formed topologies (e.g. the true topology) using the standard *RF* metric. In effect, when sequence leaves and speciation internal nodes of the offending descendant set are pruned, the resulting topology is missing a leaf, because the corresponding ancestor maps to an internal node. That node is ambiguous in its duplication vs speciation status. Nevertheless, internal nodes representing pre-duplication ancestors of the offending ancestor (and the descendant set representing it) and other ancestors of designated descendant sets can match equivalent nodes in other topologies in terms of induced partition of designated ancestors. Our modified version, *RF*\*, can account for this.

In *RF*\*, *N* represents the number of designated ancestors (descendant sets) in each compared topology, not the number of leaves. In addition to *x* and *y* we define *z* as the number of common ancestor leaves missing from *A* but not from *B* and *z*′ as the number of common ancestor leaves missing from *B* but not from *A*. The modified metric is calculated as:

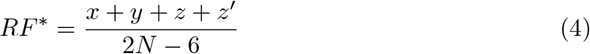

## ASPEN

ASPEN is implemented in python 2.7. The ASPEN development repository is publicly available at *https://github.com/NaegleLab/ASPEN*.

## Supporting information

**S1 Fig. Diagram of branching during ASPEN topology reconstruction**.

**S2 Fig. Log-likelihood differences between true and all-sequence topologies under JTT substitution model**.

**S3 Fig. Distributions of pairwise *RF*\* differences under 90% and 50% subsampling ensembles**.

**S4 Fig. Dropoff in ASPEN log-frequency scores across 1–precision bins**.

**S1 Appendix. Supplementary material.** Includes additional ASPEN algorithmic details and captions for figures S1 Fig through S4 Fig.

## Acknowledgments

This work was partially enabled by Center for Biological Systems Engineering. Computations were performed using the facilities of the Washington University Center for High Performance Computing, which were partially funded by NIH grants 1S10RR022984-01A1 and 1S10OD018091-01. We wish to thank Dr. Tom Ronan, Dr. Barak Cohen, Dr. Gary Stormo, Dr. Justin Fay, and Dr. Jim Havranek for the helpful discussions that shaped this work, as well as two anonymous reviewers for their helpful suggestions.

## Supplementary Material

**Figure S1:**
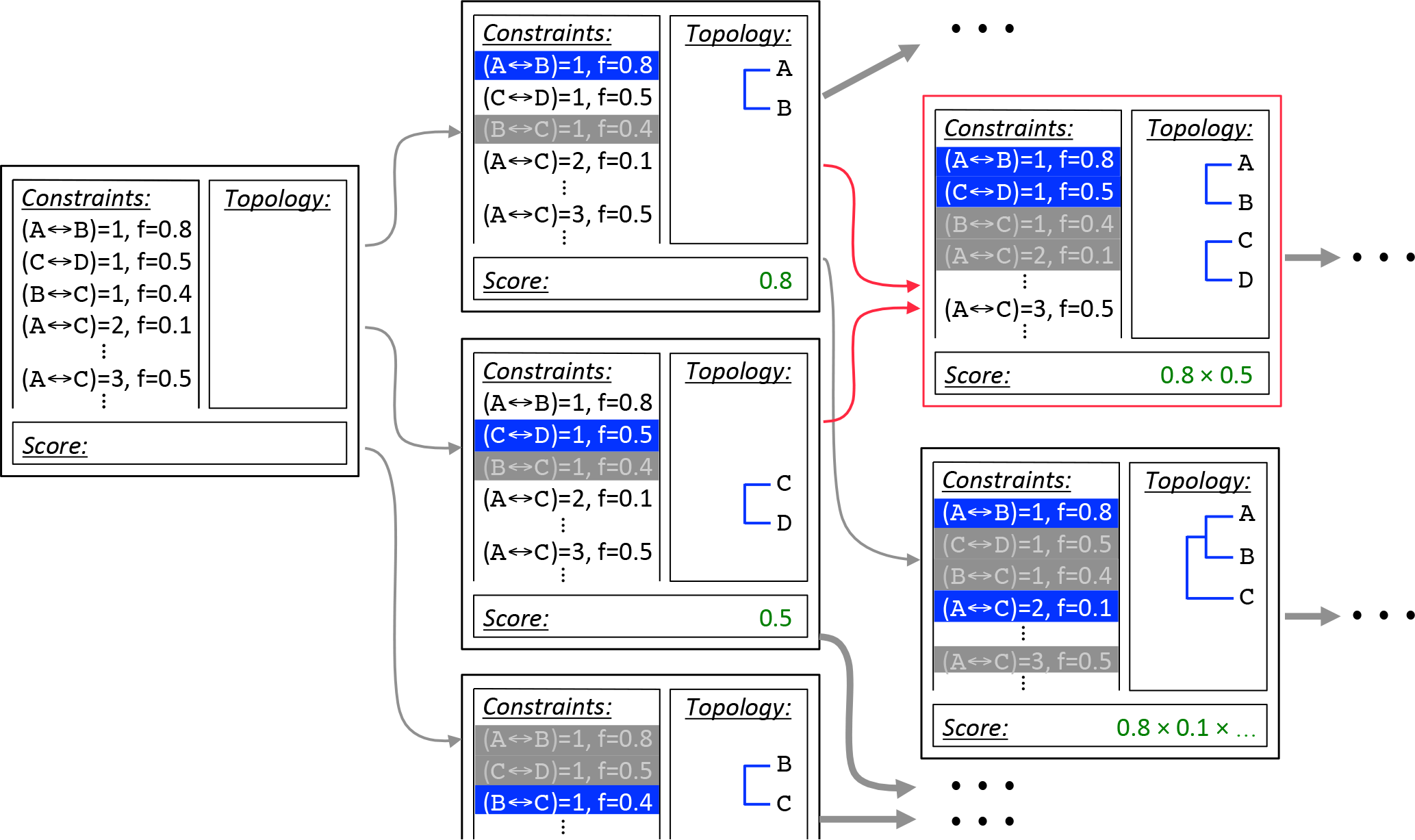
Construction begins with the empty topology assembly on the left. Every possible extension is constructed in a copy of the initial assembly: Node {*A, B*} completes path (*A ↔ B*, 1), node {*C, D*} completes path (*C ↔ D*, 1), and node {*B, C*} completes path (*B ↔ C*, 1), branching the initial assembly into three new assemblies. Path lengths completed by the introduced node and path lengths incompatible with it are marked and not revisited. Nodes {*A, B*} and {*C, D*} preclude path (*B ↔ C*, 1), while node {*B, C*} precludes paths (*A ↔ B*, 1) and (*C ↔ D*, 1). Completed paths are shown in blue, precluded paths are greyed out in the corresponding assemblies. Intermediate topology scores are calculated according to the scoring function. On the next iteration construction paths for assemblies {*A, B*} and {*C, D*} collide, indicated in red. A single copy of the resulting assembly, {*A, B*}, {*C, D*}, is retained. Assembly {*A, B*} is separately extended with node {*{A, B*}, *C*}. Additional construction sequences, indicated by ellipses, are not shown.

### Additional Algorithmic Details

This section contains additional details and examples of how assemblies are created, how they are branched, and how collisions are handled. First, the path lengths matrix is sorted into a one-dimensional list in ascending order of path length. Internal nodes are then constructed by traversing the list and joining pairs of leaves and/or previously constructed internal nodes to recapitulate encountered leaf-to-leaf path lengths. This bottom-up construction (“outside-in” for unrooted topologies) continues until all leaf nodes are connected by a single graph. In the example shown in Figure S1 construction proceeds as follows:

1. Node {*A, B*} joins leaves *A* and *B* and recapitulates path (*A ↔ B*, 1), blue.
2. Node {*C, D*} joins leaves *C* and *D* and recapitulates path (*C ↔ D*, 1), pink.
3. Node {{*C, D*}, *E*} joins leaf *E* to internal node {*C, D*} and recapitulates path (*C ↔ E*, 2), green.
4. Path (*D ↔ E*, 2), grey, was already recapitulated by the node created in the previous step, so it is skipped.
5. Node {{*A, B*}, {{*C, D*}, *E*}} joins internal nodes {*A, B*} and {*{C, D*}, *E*} and recapitulated path (*A ↔ E*, 3), orange.

- Path (*B ↔ E*, 3) and four additional paths of length 4 which appear further down in the list are also recapitulated by this node.
6. Node {{{*A, B*}, {{*C, D*}, *E*}}, *F*} joins leaf *F* to internal node {{*A, B*}, {{*C, D*}, *E*}} and recapitulates path (*A ↔ F*, 3), dashed line.

This completes the reconstruction, since all leaves are connected by the resulting topology. Path (*B ↔ F*, 3) and all subsequent paths are already recapitulated and are skipped as they are reached during list traversal. Note that with some topologies it is possible to encounter path lengths during list traversal which, at that state of construction, cannot be recapitulated by constructing an internal node. For example, if the order of paths (*A ↔ E*, 3) and (*A ↔ F*, 3) were reversed and path (*A ↔ F*, 3) was encountered first, it could not be recapitulated because internal node {{*A, B*}, {{*C, D*}, *E*}} would not yet be available to join to leaf *F*. Such path lengths are skipped and then revisited on the subsequent traversal of the list. Traversal is repeated as necessary until construction is completed. Because all path lengths are derived from a single topology, they are guaranteed to be consistent, making the construction unambiguous.

**Figure S2:**
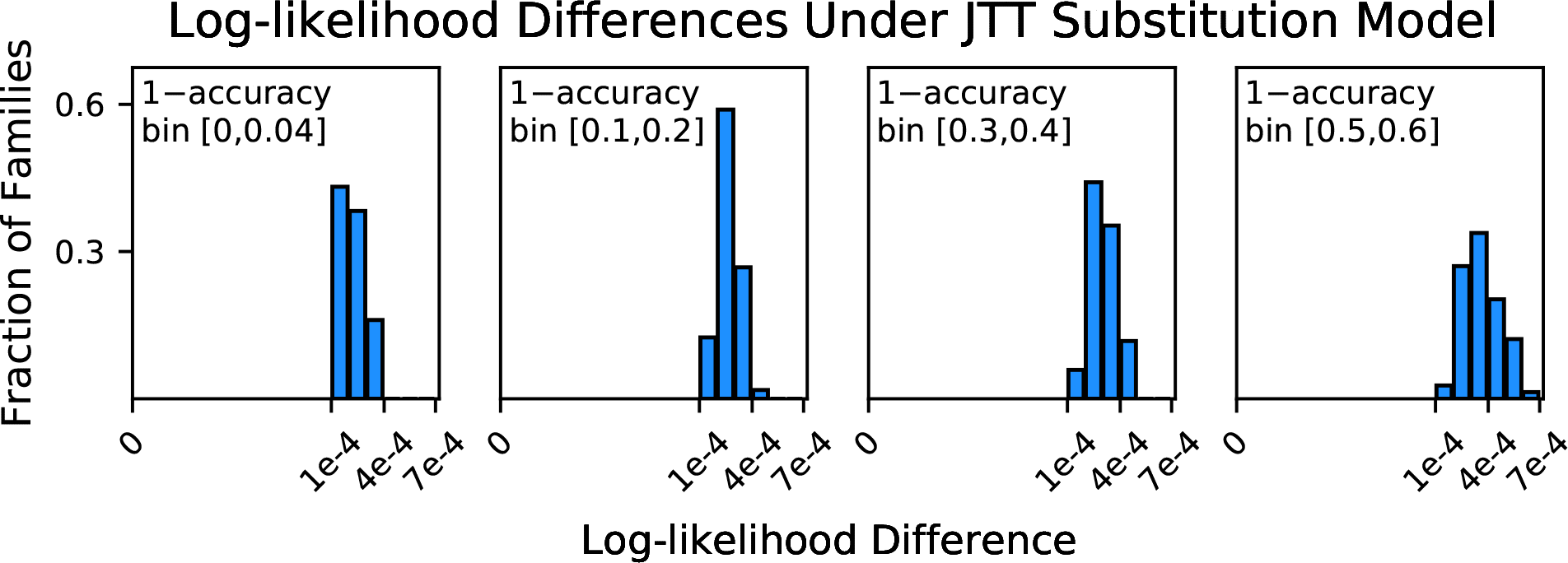
Histograms of log-likelihood differences between true and all-sequence (reference) topologies, calculated over all-sequence alignments under the JTT substitution model.

### Topology Likelihoods Under Substitution Model Used For Simulations

Evolution was simulated under the JTT substitution model, while the WAG substitution model was used for phylogenetic inferences in order to emulate the more realistic scenario where models used for inference do not precisely match actual substitution patterns. Surprisingly, not only did all true topologies have a worse likelihoods than corresponding all-sequence topologies under the inference model, WAG, the result was nearly identical under the simulation model, JTT (Figure S2).

**Figure S3:**
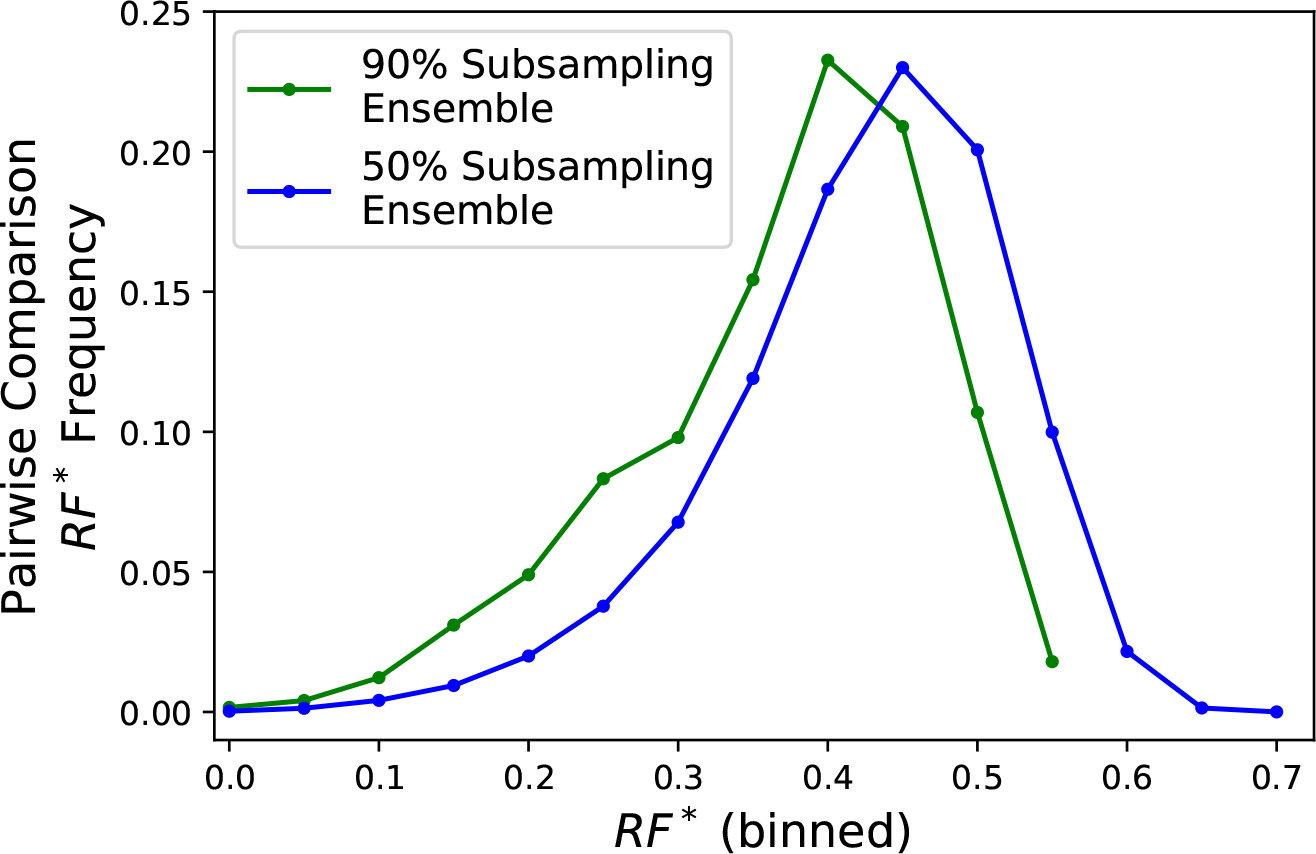
Distributions of pairwise *RF*\* differences between LacI topologies in the 90% (green) and 50% (blue) subsampling ensembles.

### Variance Among Subsampled LacI Topologies as a Function of Sampling Size

Variance of topologies created from 90% and 50% subsamples of LacI paralog sequences was measured by exhaustive pairwise comparison of all topologies in each ensemble using the modified Robinson-Foulds distance (*RF*\*, as described in manuscript). On average, topologies reconstructed from 90% subsamples are slightly more similar than those reconstructed from 50% subsamples (Figure S3), indicating more variance among the 50% subsampled ensemble.

**Figure S4:**
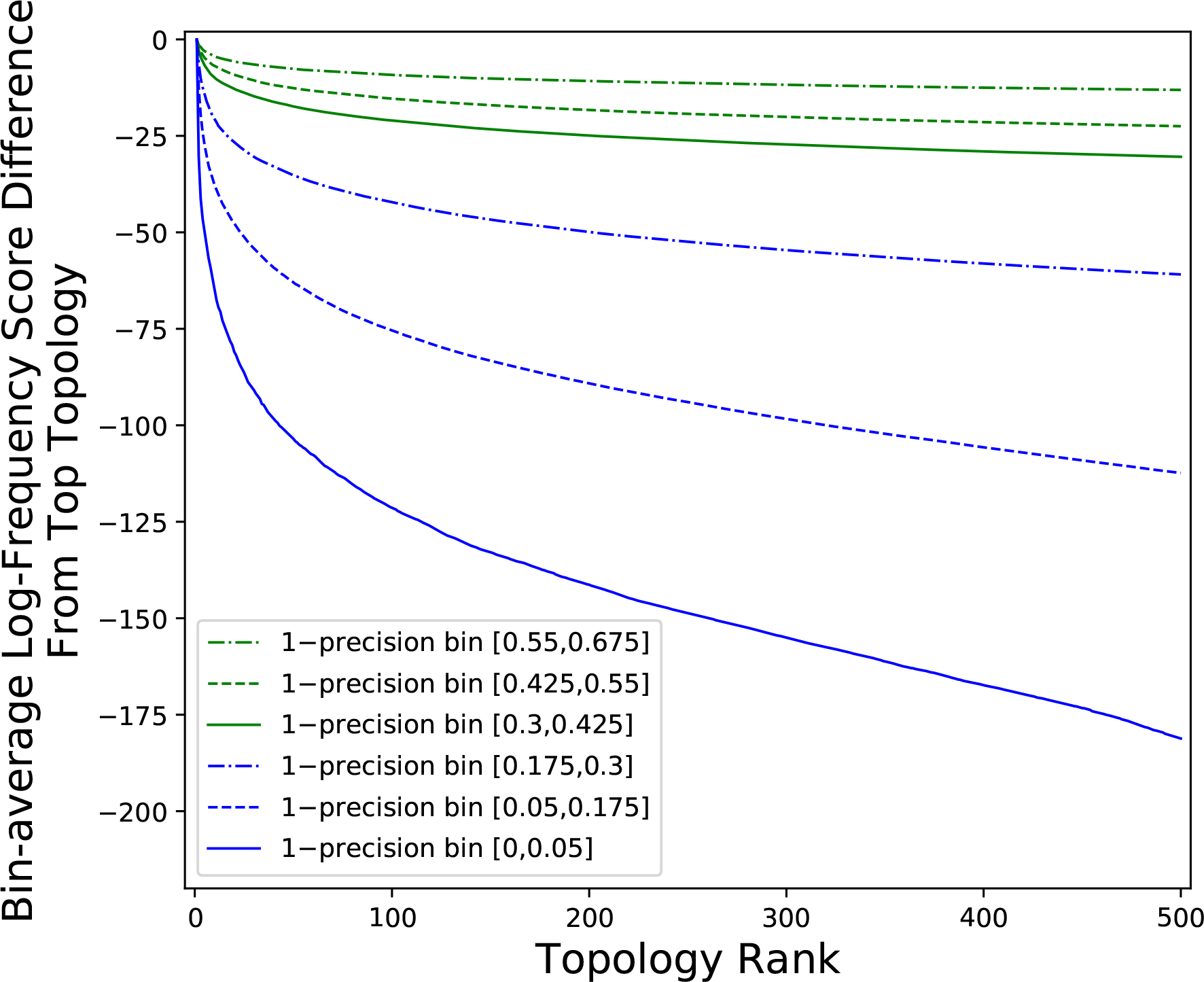
Comparison of log-frequency scores of the top 500 ASPEN topologies for simulated families across six 1–precision bins. The difference between a topology’s score and the score of the best ASPEN topology, averaged over all families in the bin, is plotted as a function of topology rank.

### Log-frequency Score Drop Off With Topology Rank Among Simulated Protein Families

The accuracy difference between the top ASPEN topology and other ASPEN topologies decreases at lower precision (Figure 4B-G, main text). Accordingly, log-frequency score differences between ASPEN topologies also decrease (Figure S4), reflecting ASPEN’s more uniform confidence in each individual topology.

### LacI Input Sequences and Topologies Reconstructed by ASPEN

1777 sequences from the LacI family, split among 23 orthosets, and the top 500 topologies reconstructed by ASPEN for the LacI family, are included in the supplemental file LacIdata.zip.

